# Female Friendships: Social Network Analysis as a tool to understand intra-group affiliation in semi-commensal lion-tailed macaques

**DOI:** 10.1101/2022.09.01.506153

**Authors:** Ashni Kumar Dhawale, Anindya Sinha

## Abstract

The lion-tailed macaque is a gregarious, rainforest-adapted species, that has, in certain locations across its natural distribution, recently begun to explore and utilise surrounding human-dominated habitats. In many primate species, exposure to novel human-use food resources and potential provisioning has previously been associated with changes in intra-group social structure, often categorised by increased aggression and, more importantly, increased affiliation, possibly as a means of reconciliation. Here we quantify the changing affiliative relationships among the female members of two groups with varying degrees of habituation, using social network analysis. We examine frequencies of pair-wise affiliation between the female members of each group, ranked within the prevailing linear hierarchy, to measure individual attributes or the local importance of individuals in a network, and group attributes or the global role of all individuals in the whole network. We found that the subordinate individuals in the less habituated group maintained a higher number of connections with group members, an expected outcome, as key affiliative behaviours such as grooming are known to be directed upwards in the hierarchy. This pattern was observed to be inverse in the highly habituated group, with dominant individuals maintaining more connections, suggesting that under the conditions of increased competition for the novel food resource, dominance rank was highly contested. In support of this theory, we also found multiple fluctuations in dominance rank over time for this highly habituated group, with nearly no fluctuations in the less habituated group. This study demonstrates that varied intensities of human presence and dependence on human-use foods have differential effects on the intra-group sociality of lion-tailed macaques.

## Introduction

Around 81% of the world’s diurnal primate species are gregarious (Sterck et al., 1997), a group-living social organisation being advantageous in terms of decreased predation risk and increased defence in cases of conflict, thus improving the chances survival for all participants (Cowlishaw, 1994; Isbell, 1994; Janson, 1992). Individuals in group-living species are, naturally, characterised as being intensely social, negotiating and maintaining social relationships with each group member, throughout their lives (Dunbar, 1988). These social relationships, which form a crucial aspect of an individual’s behavioural decisions, are likely driven by fundamental ecological factors such as the distribution of resources and risks within group home ranges (reviewed in Creel et al., 2013).

At the individual level, males and females are often considered as two separate entities, their primary behavioural decisions being ultimately governed by their physiological differences, reproductive motivations and fitness levels. In most gregarious primate species, especially among the Old World monkeys, males of the species often disperse from the natal group at sexual maturity (Greenwood, 1980). Under these circumstances, permanent group members are comprised of kin-bonded females whose social interactions are typically described through three primary modes of interaction, affiliation, aggression and dominance rank (Sterck et al., 1997). Affiliation is clearly beneficial to group members for parental care (Kappeler & van Schaik, 2002), maintaining hygiene (reviewed in Dunbar, 1991), keeping the group cohesion (Waal & Luttrell, 1982), and allyship (e.g. Brent et al., 2011), however, limits to resource acquisition creates competition among group members, leading to agonistic interactions and complex social relations in the form of dominance ranks. A dominance rank, or position of an individual in a hierarchy, denotes the measure of social power an individual holds within a group (Flack & Waal, 2004), and influences various other behavioural regimes such as foraging and movement. Dominance styles are known to vary across macaque species and range between egalitarian, where individuals are benefitted proportionate to the effort made, to despotic, where few members disproportionately benefit above others (Vehrencamp, 1983).

Even within a species, dominance styles are known to vary across subgroups and populations, depending on the nature of resources in the environment. For example, competition intensifies when food resources are clumped leading to the formation of stricter linear dominance hierarchies (Boccia et al., 1988). Most importantly, in cases when human-origin foods are accessible to wild or semi-commensal populations, an ever increasing global phenomenon, intra-group dominance patterns are seen to shift (Dhawale et al., 2020; Sinha et al., 2005). Dominance hierarchies are a key aspect for social nonhuman primates, dictating numerous behavioural choices, and even having consequences for lifetime reproductive success of an individual (Noordwijk & Schark, 1999). Predicting changes in social order, particularly in response to a changing environment, become extremely pertinent insofar as individual abilities to acquire sufficient resources, health, and fitness (Creel et al., 2013; Marmot & Sapolsky, 2014; Sapolsky, 2005) are concerned.

In this context, Social Network Analysis (SNA), a mathematical theory applied to analyse relationships between social entities (Wasserman & Faust, 1994), presents as an ideal tool. For this study, we examined a single mode of social interaction, Affiliation (Appendix 1), amongst the female members of two lion-tailed macaque groups with varied levels of human habituation, in relation to the prevailing intra-group dominance ranks.

## Study Site and Study Individuals

The study was carried out on the Valparai plateau, located in the Anamalai hill range of the southern Western Ghats, in the state of Tamil Nadu. An expanse of around 220 km^2^, the Valparai plateau is a heterogeneous landscape of rainforest fragments interspersed with tea, coffee and Eucalyptus plantations (Figure 1). With elevation ranging from 900 m to 1450 m, the plateau has a predominant native vegetation type of wet evergreen rainforest (Muthuramkumar et al., 2006). The Anamalai hills, separated from the Nilgiri hill range by the Palghat Gap, are important for the conservation of the endemic lion-tailed macaque, this being one of the eight locations where the species is present (Kumara & Singh, 2003).

**Fig 1.**
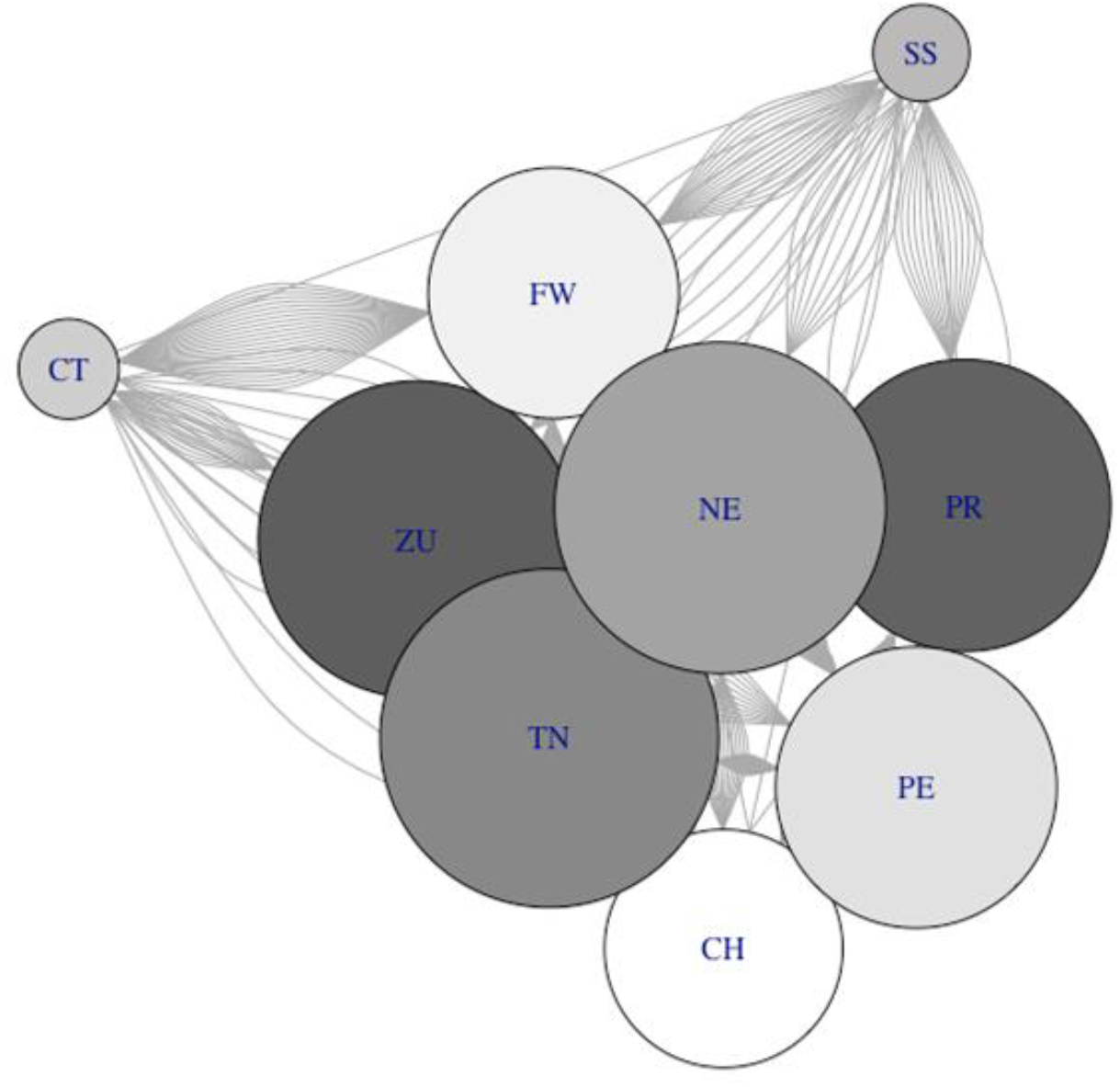
All- degree centrality for the female members of the IPH group. Node labels correspond to individual ID, colours from dark to light indicate dominance hierarchy from Subordinate to Dominant.

Since the late 1800s, extensive selective logging has led to the fragmentation and degradation of the native rainforest habitat. The lion-tailed macaque population is scattered in these remaining pockets of rainforest across the Valparai plateau. One such rainforest fragment, the Puthuthottam forest fragment, with an area of 92 ha and neighbouring the town of Valparai, is of particular importance, as it harbours a subpopulation of macaques, consisting of five groups and c.200 individuals. These groups spend significant periods of time in different anthropogenic habitats, visiting human settlements inside and around the fragment, including Valparai town, and traversing roads, orchards and plantations surrounding the fragment.

The first study group, IPH, consisted of 23 individuals, including nine adult females, one adult male, one subadult male and 12 juveniles (one to six years of age) at the start of the study period, with three infants being born subsequently. This group spent ∼80% of its time in the forest fragment, and infrequently visited human habitation. The second group, RT, consisted of 24 individuals, including eight adult females, two adult males, two subadults and 12 juveniles. This group spent nearly 90% of its time in one particular human habitation where human-use foods were openly available throughout the study period.

All the adult members of both study groups were individually identified using their distinctive features, such as facial markings, injuries or other visible abnormalities, including missing body parts or swellings/bulges.

## Methods

### Dominance Rank

For the two study groups, IPH and RT, we determined the positions of all study individual females within the prevailing linear, transitive, social dominance hierarchy through their behavioural responses during pair-wise approach-retreat interactions (A Sinha, 1998). We calculated a dominance score, namely the David’s Score or DS (Gammell et al., 2003), for each female and then categorised them as being Dominant (the top three females in the hierarchy), Subordinate (the three females with the three lowest ranks in the hierarchy) or Intermediate (the intermediate females in the hierarchy) individuals.

### Social Network

SNA, performed using igraph package (Csardi & Nepusz, 2006) in R (Team, 2013), used frequencies of Affiliation amongst the female members of each group, to represent associations in the form of a graph. Each individual is represented as a node or vertices, and a line or edges represent events of Affiliation between individuals. Measures describing individual attributes, i.e., the local importance of individuals in a network, and group attributes, i.e., the global role of individuals in light of the whole network, are defined in Table 1.

**Table 1.** The definition of each Social Network individual and group measures adapted from (Agha et al., 2020)

| MEASURES | DEFINITION |
| --- | --- |
| ***Individual measures*** |  |
| *All-degree centrality* | The number of edges which are attached to a node. |
| *Out-degree centrality* | The number of tail-ends adjacent to a node, which represent the number of initiated Affiliative behaviours with other group-members. |
| *In-degree centrality* | The number of head ends adjacent to a node, which represent the number of received Affiliations for the individual. |
| *Eigenvector centrality* | The connectivity of an individual within its network, according to the all-degree centrality of the individual and the all-degree centrality of the individuals to which it is connected. |
| *Betweenness centrality* | The number of shortest paths that pass through the considered individual. This measures the importance of the individual in connecting different subgroups in the group. |
| *Closeness centrality* | Measures how close an individual is to all individuals in the group, based on the shortest path length between that individual and all other individuals in the network. |
| ***Group measures*** |  |
| *Network density* | The number of observed edges divided by the number of all possible edges in the network. The Density value ranges between 0 and 1. |
| *Network reciprocity* | Calculates whether the relationships between two individuals are reciprocated. The Reciprocity value ranges between 0 and 1. |
| *Modularity* | The strength of the division of a network into communities. The Modularity has a value between +1 and -1, where nonzero values indicates the presence of communities within the network, while zero modularity indicates that the division of within-community edges is not different from what would be expected from the randomized network. |

### Temporal changes in Dominance Rank

We used the Elo-rating method to assess the changes in dominance ranks of the adult females in each of the study groups, over the study period. The Elo-rating method of analysis (Arpad, 1978) can overcome the shortcomings of matrix-based methods (Neumann et al., 2011), and takes into account sequential events of interactions, making it ideal to analyse temporal data. The Elo-rating scores pair-wise win-lose events by taking each participant’s previous score into consideration, and thus weighting increases or decreases in rating based on the opponent’s rating. The default Elo-rating assigned to each individual, for this study, at the start of the study period was 1000. Pair-wise approach-retreat interactions were used to examine the Elo-ratings of individuals, as they changed over the study period.

## Results

### IPH Group

The IPH group contained 9 adult females representing 9 vertices in the social networks. Of the 9 individuals, three were classified as Dominant, three as Intermediate and the remaining three as Subordinate. Individual TN, one of the Subordinate individuals, showed the highest all-degree centrality in the group, followed closely by NE, an Intermediate individual (Figure 1). The two remaining Intermediate individuals showed the least all-degree centrality. All-degree centrality was weakly correlated with dominance ranking (Spearman’s Rank Correlation r = - 0.48, p = 0.09), however, this pattern was not significant.

ZU, the bottom ranking individual in the group, showed highest out-degree centrality, followed by TN, another Subordinate individual. In-degree centrality was highest for TN, a Subordinate individual. Figure 2 depicts directional Affiliation in the IPH group, with node size indicating higher in-degree centrality. Eigenvector centrality was highest for Subordinate TN, followed by Intermediate NE, and the bottom ranking ZU. Betweenness centrality was highest for Intermediate NE, followed by a Dominant individual FW. Closeness centrality was equally high for the two most dominant individuals in the group CH and FW, an Intermediate NE and two subordinate individuals TN and ZU. The two remaining Intermediate females, CT and SS remained consistently low on all centrality measures.

**Fig 2.**
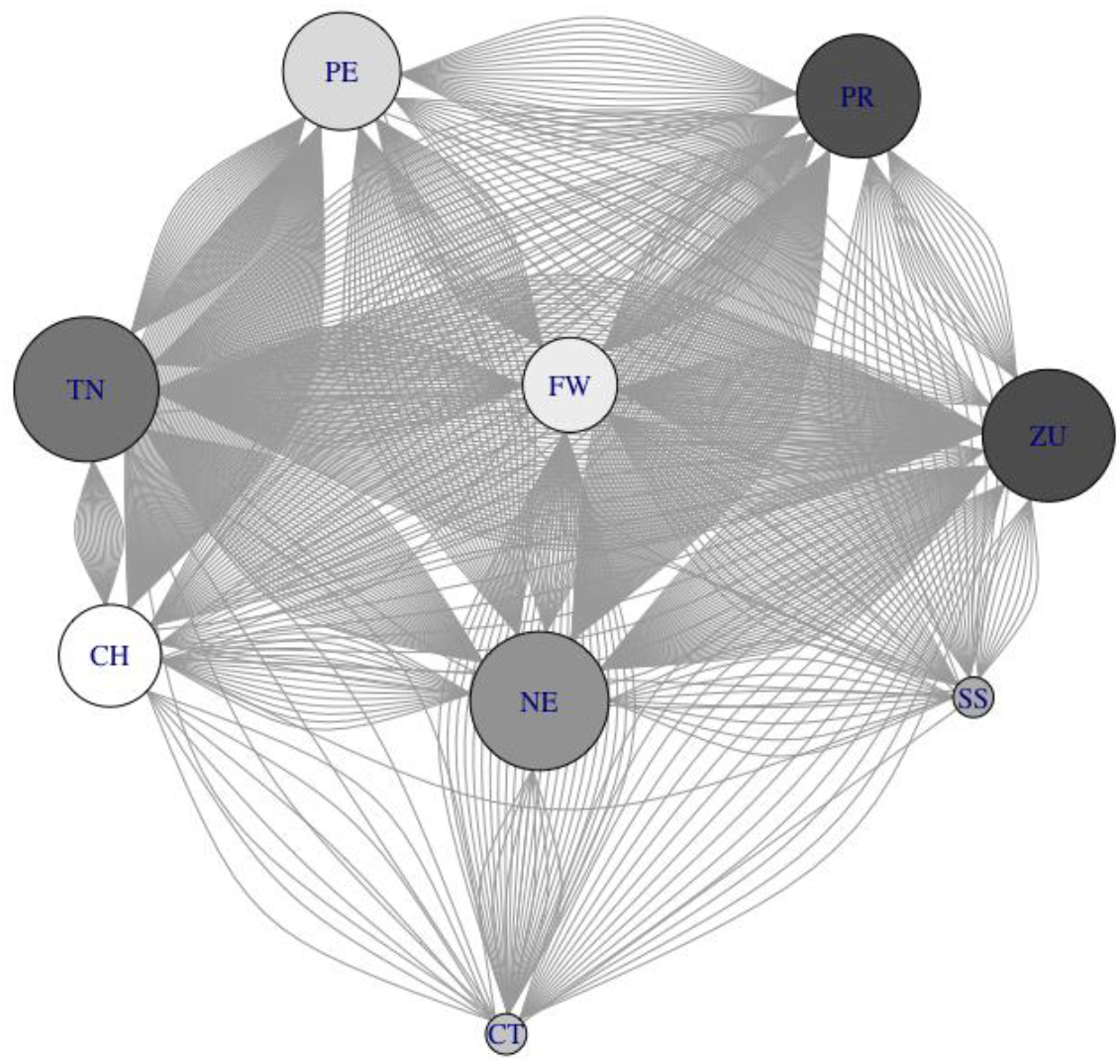
In-degree centrality for the female members of the IPH group. Node labels correspond to individual ID, colours from dark to light indicate dominance hierarchy from Subordinate to Dominant.

The network density for the IPH group was measured at 0.79 and network reciprocity was measured at 0.70. Four communities were detected in the IPH group (Figure 3), and the network modularity was measured at 0.039.

**Fig 3.**
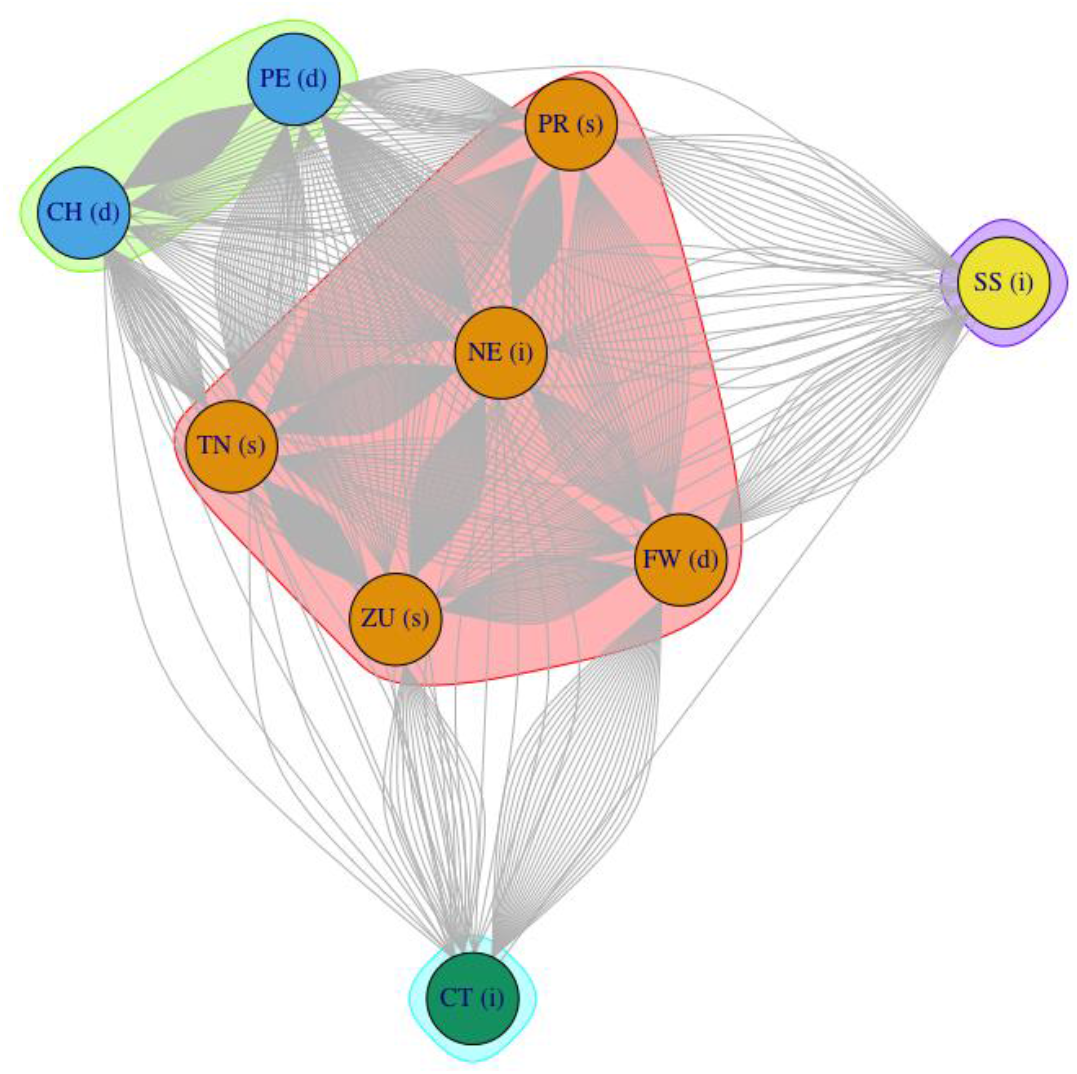
Communities, with delineating polygons, detected in the IPH group

### RT Group

The RT group consisted of 8 adult females, representing 8 vertices in the social networks. Of these, three were classified as Dominant, two were classified as Intermediate and three were classified as Subordinate. LN, a Dominant individual showed the highest all-degree centrality, followed by an intermediate individual WE (Figure 4). The most Dominant individual and the bottom ranking individual showed the least all-degree centrality. There was no observed relationship between dominance rank and all-degree centrality (Spearman’s Rank Correlation r = -0.095, p = 0.84).

**Fig 4.**
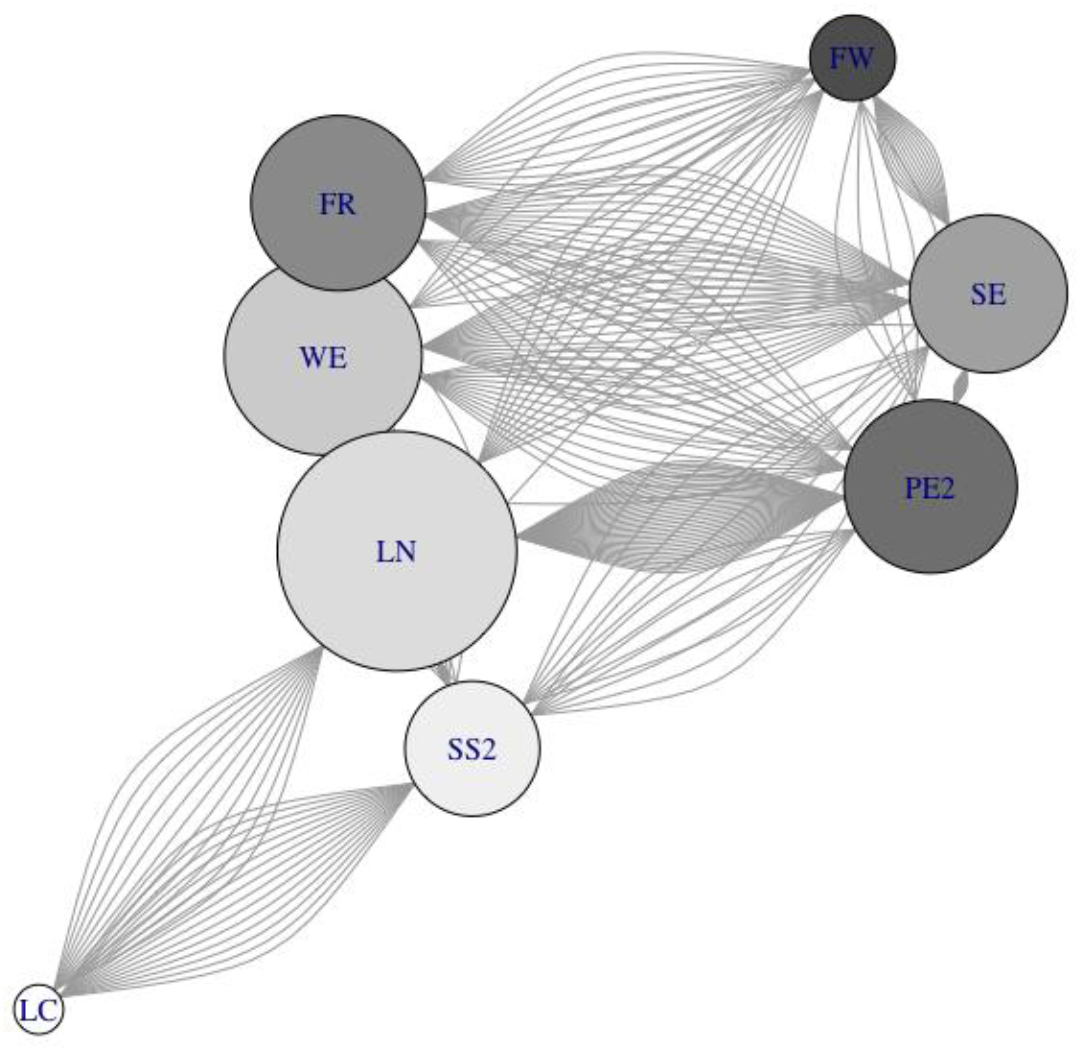
All- degree centrality for the female members of the RT group. Node labels correspond to individual ID, colours from dark to light indicate dominance hierarchy from Subordinate to Dominant.

PE2, a Subordinate female showed the highest out-degree centrality, followed by FR, another Subordinate individual, while a Dominant female, LC, showed the least out-degree centrality. In-degree centrality was highest for LN, a Dominant individual, followed by Intermediate individual WE, and was least for the bottom-ranking female (Figure 5). LN also had the highest eigenvector centrality, closely followed by Intermediate WE and Subordinate PE2. For betweenness centrality, once again, LN showed the highest measures, followed by another Dominant individual, SS2. Four individuals received equal scores for closeness centrality, Dominant LN, Intermediate WE, and Subordinates FR and PE2. The top-ranking female, LC, and the bottom ranking female, FW, consistently scored low in all measures of centrality.

**Fig 5.**
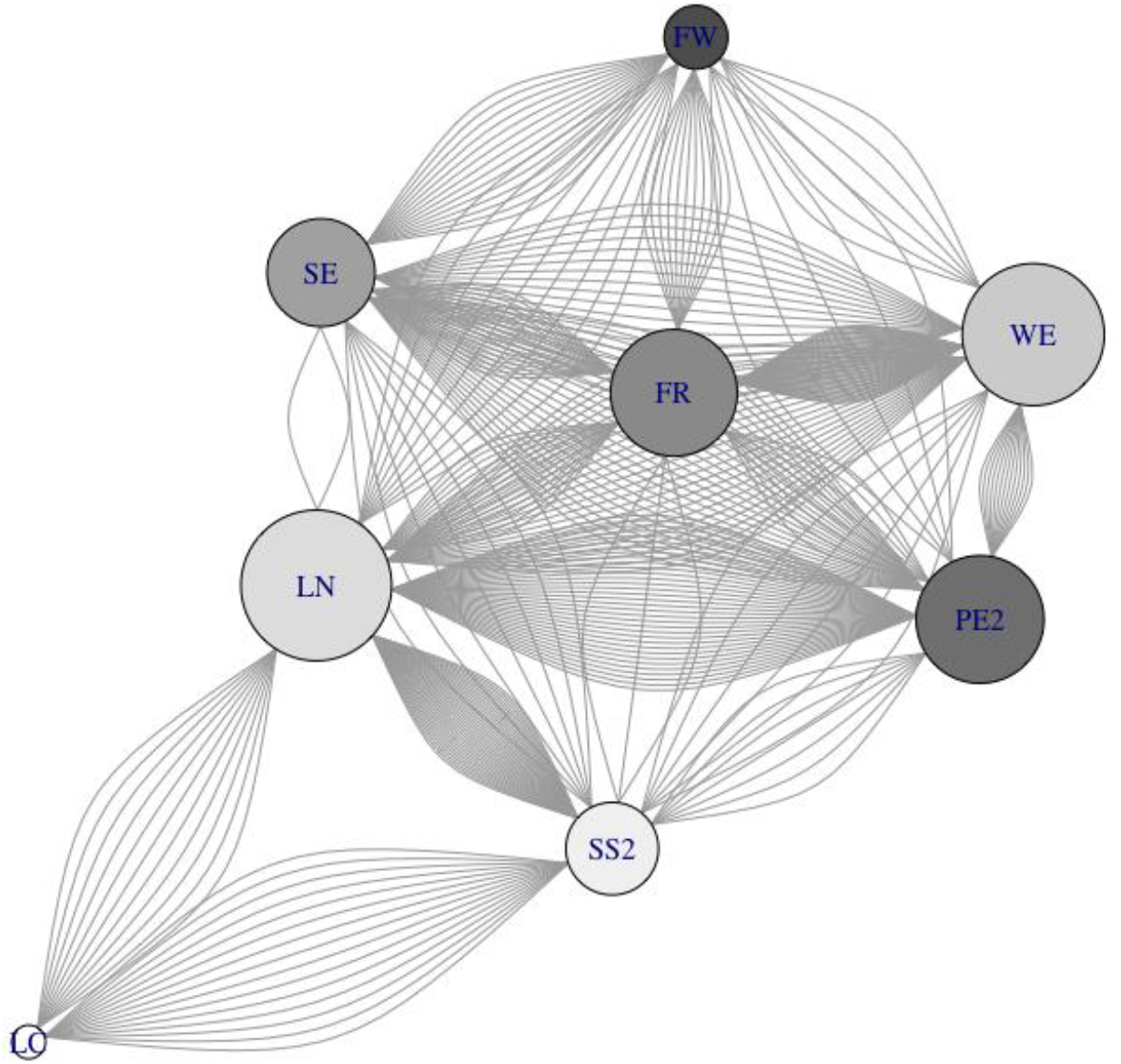
In-degree centrality for the female members of the RT group. Node labels correspond to individual ID, colours from dark to light indicate dominance hierarchy from Subordinate to Dominant.

The network density for the RT group was measured at 0.64 and network reciprocity was measured at 0.43. Four communities were detected in the RT group (Figure 6), and the network modularity was measured at 0.062.

**Fig 6.**
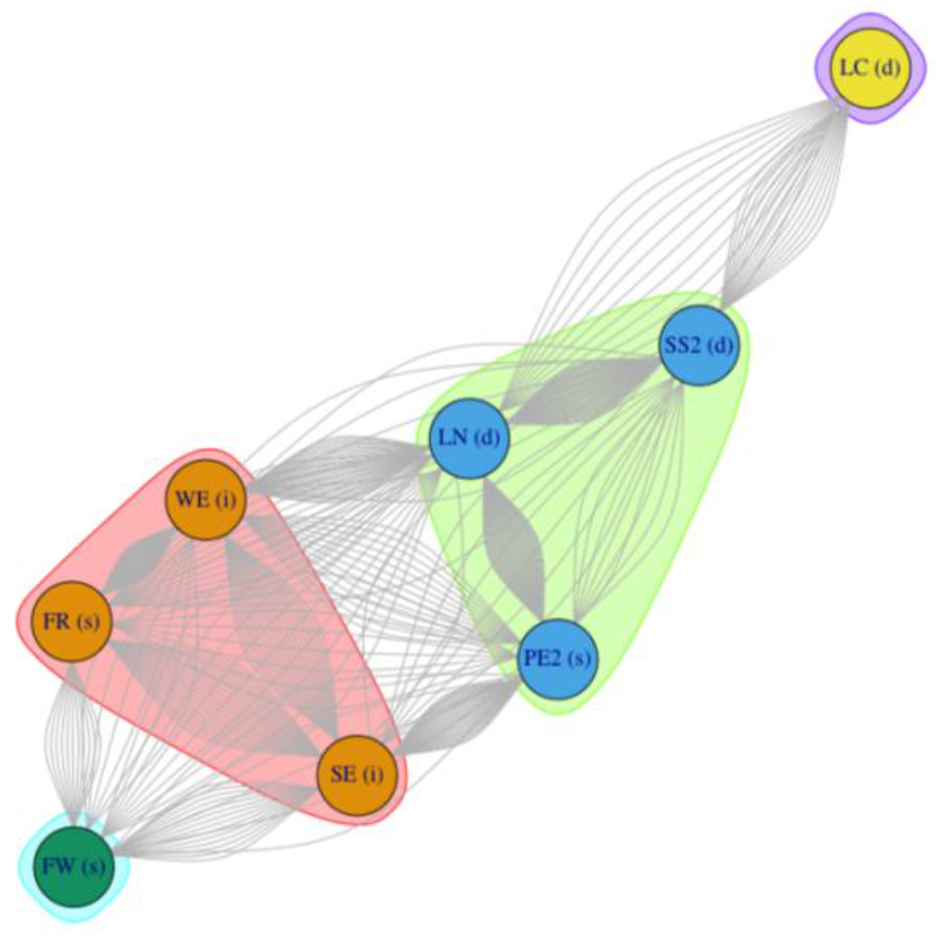
Communities, with delineating polygons, detected in the RT group

The Elo-ratings for the IPH Group ranged between 800 to >1200 (Figure 7), whereas the ratings for the RT Group ranged between 700-1400 (Figure 8). The RT Group also exhibited high fluctuations in individual ranking over the first six months of the study period, which was not the case for the IPH Group.

**Fig 7.**
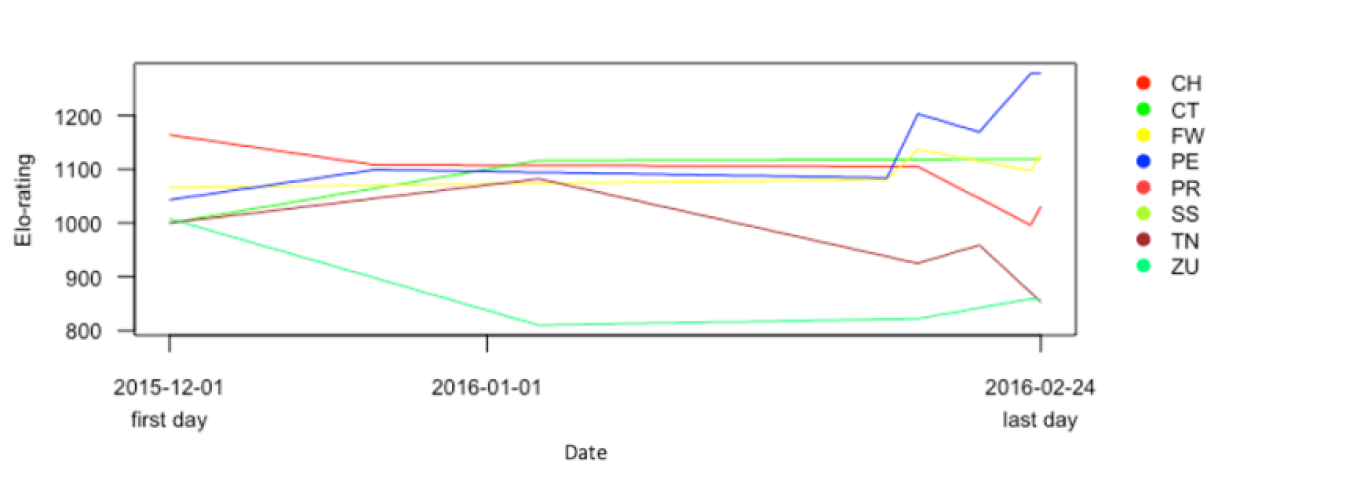
Change in dominance rank for IPH Group over the study period

**Fig 8.**
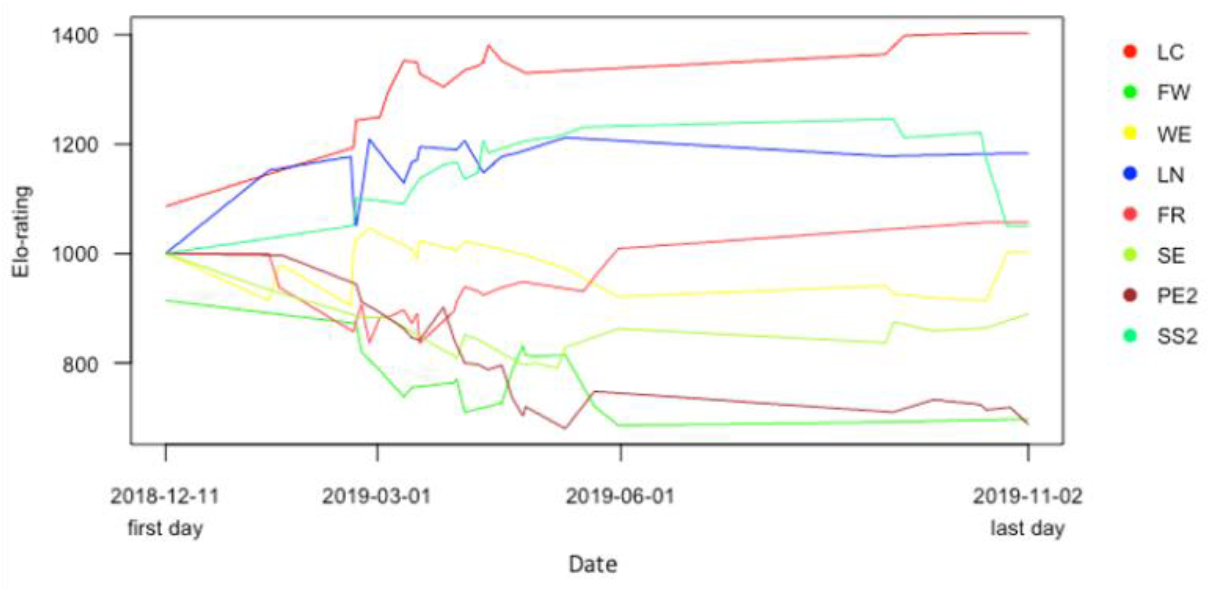
Change in dominance rank for RT Group over the study period

## Discussion

Patterns of increased affiliation, or reconciliation, are commonly observed in numerous macaque species under provisioning regimes (Ram et al., 2003; Gumert, 2010; Matsuzawa & Yamagiwa, 2010; Dhawale et al., 2020), providing crucial insights into the changing social structures along a gradient of human influence. For example, in macaque populations that are exposed to human-use foods, high ranking individuals have disproportionately higher access to food sources, consequently restricting certain individuals from reaping these benefits (Marty et al., 2020). Additionally, the presence of humans in the vicinity of macaques is known to directly or indirectly impact social behaviours, leading to decreased overall group cohesion, which has major implications for long-term survival in anthropogenic habitats (Marty et al., 2019).

Our study examines the social interactions among the female members of two lion-tailed macaque groups with varying levels of human habituation. The IPH Group visited human habitation infrequently and used only certain areas with low human presence, such as the Iyerpadi Garden Hospital. Conversely, the RT Group spent >90% of its time in a high-density human settlement, Rottikadai, and was relatively more habituated to human presence. Both groups were similar in group size and composition with the IPH Group having nine adult females and the RT Group consisting of eight adult females.

Our investigations on the social structure of adult females in the IPH and RT Groups through SNA revealed certain noteworthy differences. Degrees of centrality in the IPH Group, measuring the connections maintained by individuals in the group, were highest for Subordinate individuals, whereas the most influential member, or the one that showed highest mediations between subgroups in the group, was in Intermediate individual. Affiliative behaviours, such as grooming, are known to be directed up the hierarchy, this being demonstrated in the lion-tailed macaque (Mridula Singh et al., 2006b) and other macaque species (Furuichi, 1984; Cooper & Bernstein, 2000), thus, it was unsurprising that the Subordinate individuals in the IPH group maintained the most connections. The group also showed a relatively high network density, indicating strong social cohesion, also corroborated by the closeness measure for which five out of the nine individuals scored equally. Reciprocity of affiliation was also measured as relatively high. In detecting community structure, it is worth noting that the SNA analysis is not robust against smaller resolution local community detections, thus the modularity measures are exceptionally low, however, four distinct subgroups were detected in the IPH Group. These subgroups were divided based on both age and dominance rank, with the largest subgroup containing all Subordinate individuals, one Intermediate individual and one Dominant individual, the second containing only two Dominant females, and the remaining two were socially isolated forming their own independent subgroups. Interestingly, the two socially isolated individuals, both belonging to the Intermediate rank, which scored consistently low on all individual measures, were the oldest and the second-from-oldest members of the group. The age of the individuals was ascertained by visual inspection (National Research Council, 1981); the two oldest females had a scrawny physique, gnarled fingers and joint swellings, these being more advanced in the older of the two. In this case, age, rather than dominance rank appeared to be the predominant driver for the scarce engagement in social interactions with other group members. This pattern, of older females selectively withdrawing from social interactions, has, in fact, been reported in two other macaque species, the stump-tailed macaque and the Japanese macaque, being explained by lowered benefits for aged members of a group, and reduced energy for complex negotiations (Hauser & Tyrrell, 1984).

The RT Group showed a slightly varied social structure. In terms of degree centralities, a Dominant individual LN maintained most connections, scored highest as mediator between subgroups and received the most affiliative interactions from group members. FR, a Subordinate individual gave the most affiliation to others in the group, and LC, the top-ranking individual gave the least affiliation. While this pattern also indicates an upward direction of affiliation, where a Subordinate initiated associations most and a Dominant initiated them least, it is interesting to note that a single Dominant individual interacted with many individuals in the group and received most affiliation from the others, a role that was assumed by an Intermediate individual in the IPH Group. The RT Group also showed lower network density and reciprocity than the IPH Group, with only half the group members showing a high closeness score. Local communities in the RT Group, while also having four distinct subgroups, seemed to be divided based on dominance rank entirely, and indicated a tiered structure between low-ranking and high-ranking individuals. The first subgroup contained a Subordinate individual associating most frequently with two Intermediate individuals, the second contained a Subordinate individual associating most frequently with two Dominant individuals and the top-ranking and bottom-ranking individuals formed two socially isolated independent subgroups. While it was relatively unexpected to find a Subordinate individual associating directly with Dominant individuals, this individual appeared to be especially bold and persistent. Additionally, we observed numerous instances of aggression shown towards the bottom-ranking individual by the top-ranking individual, which seems to have led to her social isolation. We believe that under the conditions of increased competition for this habituated group, dominance rank was conceivably highly contested, and most individuals, despite being in the upper ranks, needed to maintain associations to protect their positions in the hierarchy, consequently resulting in lowered reciprocity within dominance categories and reduced overall social cohesion.

The change in dominance rank for individuals over the study period varied drastically for the two study groups. We observed very few instances of fluctuations in individuals’ dominance rank in the IPH Group, whereas for the RT Group, there were frequent fluctuations in the first six-month period, but these seemed to stabilise in the second six-month period. The unusual fluctuations in the RT Group can perhaps be explained by certain circumstances under which this group formed. Unlike the IPH Group, the RT Group was newly formed, resulting from a previous fission of two other groups in the populations, followed by the fusion of a subset of members from each group. These individuals were likely interacting for the first time with unrelated conspecifics, and subsequently clamouring to acquire positions in the dominance hierarchy. We believe the fluctuations in the initial six-month period reflected this process, which then stabilised over time. What is noteworthy, however, is that in the case of both groups, individuals rarely wavered outside their dominance categories, suggesting that certain individuals, likely having certain traits, assumed a Dominant status from the very beginning and were able to maintain it throughout the study period.

Our study demonstrates that varied intensities of human presence and dependence on human-use foods have differential effects on the intra-group sociality of lion-tailed macaques. Our observations follow patterns previously observed in other macaque species in anthropogenic habitats, and throw light on the importance of understanding social structures in response to rapidly changing environments. In this context, the observed shifts in social regimes due to the apparent social stress created by the human-dominated habitats in the study area highlight the increased vulnerability of habituated lion-tailed macaques, having substantial implications on the long-term survival ability of the population.

## Notes

### Competing Interest Statement

The authors have declared no competing interest.

